# *Atp6v1b2* Plays Important Roles in the Early Development of Hearing, the Pectoral Fin, the Cardiovascular System, and the Swim Bladder of the Zebrafish, Supporting a Role for the Gene in Syndromic Hearing Loss

**DOI:** 10.1101/024240

**Authors:** Yongyi Yuan, Xue Gao, Feng Xin, Pu Dai

**Affiliations:** Department of Otolaryngology, Chinese PLA General Hospital, Beijing, 100853, People’s Republic of China; Department of Otolaryngology, Hainan Branch of PLA General Hospital, Sanya, 572000, People’s Republic of China

**Keywords:** *atp6v1b2*, development, syndromic hearing loss, knockdown

## Abstract

**Background:** The *ATP6V1B2* gene plays a critical role in the auditory system, and a mutation in this gene is one genetic cause of DDOD syndrome (Dominant deafness-onychodystrophy syndrome, MIM 124480) and ZLS (Zimmermann-Laband syndrome, MIM 135500). However, whether and how *ATP6V1B2* is involved in the development of other organs remains unknown. In the present study, we explored the effect of *atp6v1b2* knockdown on early zebrafish development and verified that this gene plays a role in syndromic hearing loss.

**Methods:** Three morpholinos (two splice-blocking, one translation-blocking) and the *atp6v1b2* c.1516 C>T plasmid were used to knockdown or overexpress atp6v1b2 in zebrafish after microinjection of fertilised embryos. Control and *atp6v1b2* embryo morphants were evaluated 6 days post-fertilisation in terms of motility, apoptosis, pectoral fin development, and hair cell number.

**Results:** *Atp6v1b2*-knockdown zebrafish exhibited decreased body length, pericardial oedema, hair cell loss, a non-inflated swim bladder, and shorter pectoral fins; the first three phenotypes were also evident in fish overexpressing the gene.

**Conclusions:** *Atp6v1b2* plays important roles in the development of hearing, the pectoral fin, the cardiovascular system, and the swim bladder, thereby supporting a role for this gene in syndromic hearing loss.

## Introduction

Hearing loss is one of the most common sensory disorders. In 2013, the World Health Organization (WHO) reported that over 360 million people suffered from this condition (WHO 2013). In newborns, the incidence of profound hearing loss is 1:1,000 (Morton 1991). About 50–60% of cases have genetic etiologies, 25% are caused by environmental factors, and the remaining 25% are of unknown origin (Morton 1991; Schrijver 2004). Genetic hearing loss may develop alone or as part of a syndrome. About 70% of all genetic hearing loss is nonsyndromic.

Currently, more than 1,000 syndromes featuring hearing loss are recognised in the London Dysmorphology Database (Tseng and Lalwani 2000). Morbidity and mortality associated with such loss varies with the extent of involvement of non-auditory systems. For example, children with Jervell and Lange-Nielsen syndromes are at risk of syncope, arrhythmia, and sudden death (Chiang and Roden 2000). Children with Usher syndrome develop hearing loss accompanied by vestibular and visual impairment. Usher syndrome is the etiology of most combinations of deafness and blindness. Such dual sensory impairment has enormous implications in terms of both communication and education (Friedman et al. 2011). Glomerulonephrosis associated with Alport syndrome can trigger kidney failure, necessitating a kidney transplant (Gumber et al. 2012). Thus, knowledge of the molecular pathogenesis of syndromic hearing loss can guide clinical and pre-implantation genetic diagnoses, facilitating early intervention, genetic counselling, risk assessment, and the choice to bear or not bear children.

Dominant deafness-onychodystrophy (DDOD) syndrome (MIM 124480) is a type of ectodermal dysplasia characterised principally by congenital sensorineural hearing loss accompanied by dystrophic or absent nails. In some individuals, conical and hypoplastic teeth may also develop. The principal difference between DDOD and DOORS (deafness, onychodystrophy, osteodystrophy, intellectual disability, and seizures, MIM 220500) syndrome is that DOORS is associated with intellectual disability and seizures. Mutations in *TBC1D24* were recently shown to cause DOORS syndrome (Campeau et al. 2014b; Campeau et al. 2014a). To date, 10 families (of varying ethnicity) with DDOD syndrome have been described (Feinmesser and Zelig 1961; Goodman et al. 1969; Kondoh et al. 1999; Moghadam and Statten 1972; Robinson GC 1962; Vind-Kezunovic and Torring 2013; White and Fahey 2011; Yuan et al. 2014). We previously found that the *ATP6V1B2* c.1516 C>T mutation co-segregated with certain phenotypes of DDOD syndrome, and that the gene plays a critical role in the auditory system. However, whether and how *ATP6V1B2* is involved in the development of other ectodermal organs such as nails, toes, and the phalanx, remain unknown.

The zebrafish (*Danio rerio*) embryo is a valuable model of vertebrate development, affording many practical advantages including external fertilisation, an optically clear chorion, a translucent embryo, rapid development, and accessibility of early developmental stages. Thus, the zebrafish is an excellent model of vertebrate embryonic development and the genetic analysis of such development. In the present study, we explored the effect of *atp6v1b2* knockdown on the early development of zebrafish, and verified that this gene plays a role in syndromic hearing loss.

## Materials and Methods

### Zebrafish care and maintenance

Adult wild-type AB strain zebrafish were raised and maintained under standard conditions (M. 1993). Embryos were staged as previously described (Kimmel et al. 1995). The Animal Care and Use Committees of the Chinese PLA General Hospital and the Shanghai Biomodel Organism Science & Technology Development Co. Ltd approved all of the animal procedures used, which were consistent with those of the American Veterinary Medical Association Panel on Euthanasia.

### Zebrafish microinjection

Gene Tools LLC (http://www.gene-tools.com/) designed the morpholinos (MOs) used. Antisense MOs were microinjected into fertilised one-cell-stage embryos using a standard protocol (Nasevicius and Ekker 2000). The sequences of the *atp6v1b2* translation-blocking and splice-blocking MOs were 5’- TCACCATTCCCCTAAGAGCCTTCAT-3′ (ATG-MO), 5′- ATGTTTTTTCTAAATCTCACCCAGC-3′ (E4I4-MO), and 5′- CCTACCATTGTTACTTACCCTGGGA-3′ (E13I13-MO). In total, 4 ng each of E4I4-MO and ATG-MO were injected per embryo, in addition to 8 ng E13I13-MO. Total RNA was extracted from groups of 80–100 embryos using the Trizol reagent (Invitrogen) according to the manufacturer’s instructions. RNA was reverse-transcribed using the PrimeScript RT reagent Kit with the gDNA Eraser (Takara). Primers spanning *atp6v1b2* exon 2 (forward primer: 5′- AAACTGTGGCTGGTGTTAATG-3’) and exon 5 (reverse primer: 5′-GGCTGTCCCATAATGTCTAAGT-3’) were used in RT-PCR to confirm that the E4I4-MO was active. Primers spanning *atp6v1b2* exon 12 (forward primer: 5′-GGTGAGGGAATGACACGTAAA-3′) and exon 14 (reverse primer: 5′-TCACGGTTGATTGAGAGACAAA-3′) were used in RT-PCR to confirm the efficacy of E13I13-MO.

In the overexpression assay, fertilised one-cell embryos were injected with 200 pg pcDNA3 containing human mutant *ATP6V1B2* cDNA (concentration: 100 ng/μL). Control and *atp6v1b2* morphant embryos were allowed to develop to 6 days post-fertilisation (dpf) prior to evaluation of motility, apoptosis, pectoral fin development, and hair cell number.

### Behavioral assays

Larvae from several 48 h clutches were pooled. Selected fish were manually dechorionated at least 3 h before the experiment. To evaluate the escape response, fish were touched with the tip of a fine needle at least twice, at the dorsal tips of the tail or trunk. An escape response whereby a fish did not move a distance of at least three body lengths was considered reduced. One experiment was recorded and video-processed.

***Acridine orange staining to detect apoptosis***

Wild-type control embryos and embryos injected with 4 ng *atp6v1b2*-e4i4-MO were immersed in 5 μg/mL acridine orange (AO) (acridinium chloride hemi[zinc chloride salt], Sigma) in fish water for 60 min at 32 h post-fertilisation (hpf). Next, the fish were thoroughly rinsed three times in fish water (5 min/wash), and anaesthetised with 0.016% (w/v) tricaine methanesulfonate (TMS) (MS-222, Sigma). Then, the fish were oriented laterally, and mounted (using methylcellulose) in the depressions of appropriate slides prior to fluorescence microscopy.

### DASPEI staining of lateral line hair cells

Zebrafish embryos microinjected at 5 dpf were immersed in 1 mM DASPEI (2-(4-(dimethylamino)styryl)-N-ethylpyridinium iodide, Sigma) in fish water for 1 h, rinsed thoroughly three times in fish water (5 min/wash), anaesthetised with 0.016% (w/v) TMA, oriented laterally (anterior, left; posterior, right; dorsal, top), and mounted (using methylcellulose) in the depressions of appropriate slides prior to fluorescence microscopy.

### Image acquisition

Embryos and larvae were examined using a Nikon SMZ 1500 fluorescence microscope, and photographed with a digital camera. Subsets of images were adjusted in terms of level, brightness, contrast, hue, and saturation using Adobe Photoshop 7.0, to optimally visualise expression patterns. Quantitative image analyses were performed with the aid of image-based morphometric (NIS-Elements D3.1, Japan) and ImageJ software (United States National Institutes of Health; http://rsbweb.nih.gov/ij/). Inverted fluorescent images were processed. Positive signals were quantified (particle numbers) with ImageJ software. Ten fish were subjected to each treatment, and signal numbers/fish were calculated by data averaging.

### Statistical analysis

All of the data are presented as means ± SEMs. Statistical analysis and graphical data representation were performed using GraphPad Prism 5.0 (GraphPad Software, Inc., La Jolla, CA). Statistical significance was assessed using Student’s *t*-test, ANOVA, or the χ^2^ test, as appropriate. Statistical significance is indicated by * (P < 0.05) and *** (P < 0.0001).

## Results

### atp6v1b2 knockdown causes developmental defects in zebrafish

Zebrafish injected with *atp6v1b2* E4I4-MO (4 ng), ATG-MO (4 ng), or E13I13-MO (8 ng per embryo) all showed altered phenotypes. The proportions of embryos exhibiting defects were 65.41% (104/159), 75.90% (189/249), and 47.30% (70/148), respectively. Compared to wild-type, *atp6v1b2-*knockdown zebrafish hosting E4I4-MO and ATG-MO had a shorter body length, severe pericardial oedema, and a non-inflated swim bladder (Figs. 1, S1, S2), whereas those knocked down with E13I13-MO only presented with the latter two defects (Fig. S3). Both *atp6v1b2*-e4i4-MO and *atp6v1b2*-e13i13-MO were active, as confirmed by RT-PCR (Figs. S4, S5). At 32 hpf and 5 dpf, zebrafish embryos injected with 4 ng *atp6v1b2*-e4i4-MO had a significantly decreased relative body length (X10^2^ μm) (20.40±0.4322 vs. 22.46±0.1370; 32.89±1.501 vs. 37.74±0.5213, respectively; P<0.0001). At 32 hpf, zebrafish embryos injected with 4 ng *atp6v1b2*-ATG-MO also had a significantly decreased relative body length (X10^2^μm) (20.32±0.9046 vs. 22.46±0.1370; P<0.0001). The stereotypic escape response was dulled in *atp6v1b2*-e4i4-MO-injected 48 hpf larvae (Supplementary Movie). No significant organ-specific apoptosis was evident in either wild-type control or *atp6v1b2* morphants.

**Fig. 1.**
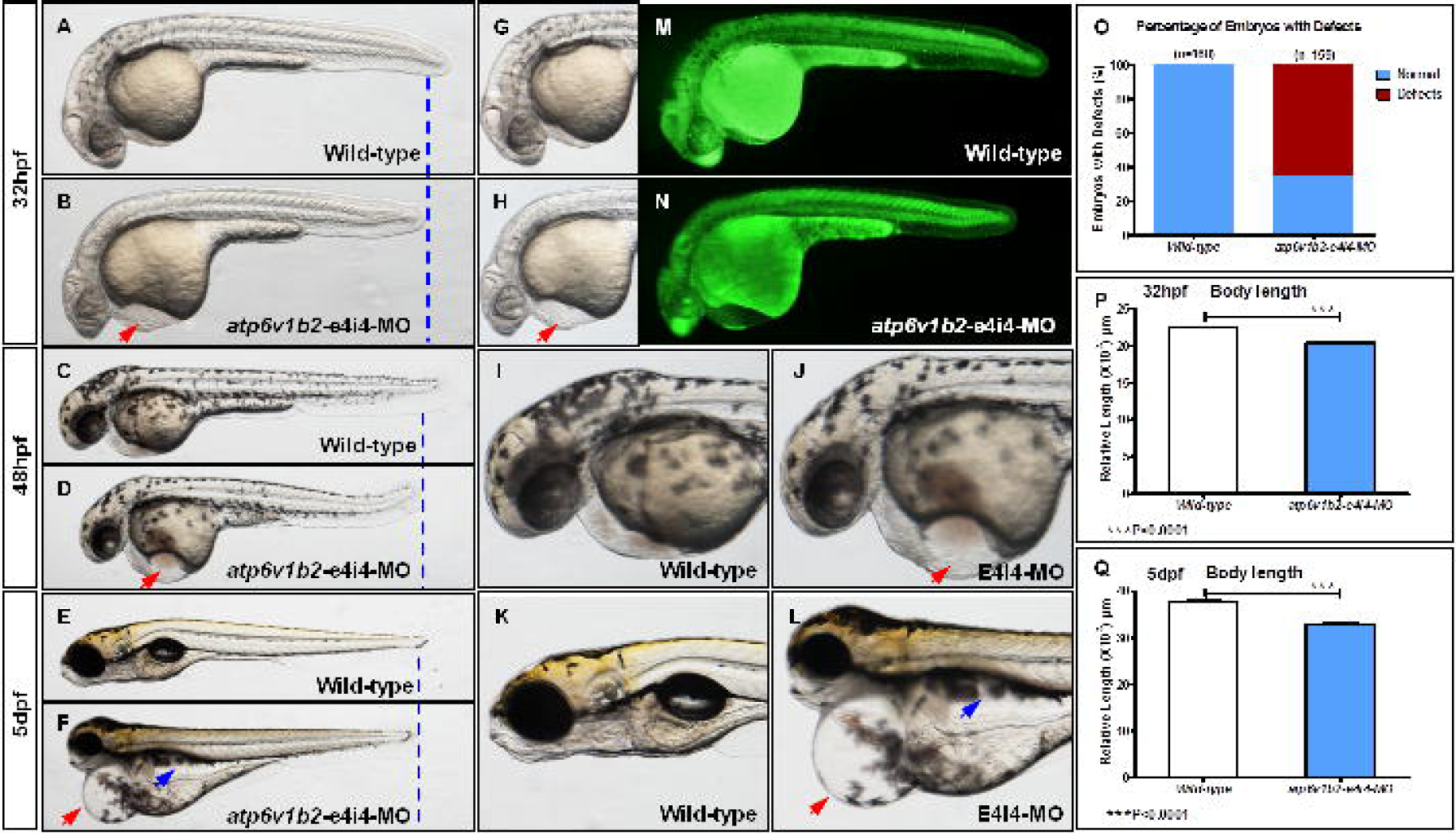
Aberrant *atp6v1b2* triggers developmental defects in zebrafish. (A-F) Gross morphologies of wild-type (WT) embryos or embryos injected with 4 ng *atp6v1b2*-e4i4-MO, at 32 hpf, 48 hpf, and 5 dpf (B, D, F, H, J, L). Compared to WT embryos, *atp6v1b2* knockdown decreased body length (blue dotted line), caused severe pericardial oedema (red arrow), and inhibited inflation of the swim bladder (blue arrow) (Figs. S1–S3). (M–N) Acridine orange staining did not evidence any significant organ-specific apoptosis in either the WT control or *atp6v1b2* morphants. The stereotypical escape response was abnormal in *atp6v1b2*-e4i4-MO-injected larvae at 48 hpf (Supplementary Movie). The bar graphs in Panels O through Q show the percentages and body lengths of embryos with developmental defects. RT-PCR confirmed that the *atp6v1b2*-e4i4-MO was effective (Figs. S4, S5). hpf, h post-fertilisation; dpf, days post-fertilisation.

### atp6v1b2 knockdown disrupts pectoral fin development

Fins and limbs are thought to be homologous organs; the developmental mechanisms are similar. The development of paired zebrafish fins (e.g., the pectoral fins) and tetrapod forelimbs and hindlimbs exhibit striking similarities at the molecular level (Gehrke et al. 2015; Ma et al. 2009; Mercader 2007; Neto et al. 2012; Pi-Roig et al. 2014; Rothschild et al. 2009; van der Velden et al. 2012; Yano et al. 2012). In uninjected wild-type control zebrafish, the pectoral fins extended out from the body wall and were of normal length at 6 dpf. In 6 dpf *atp6v1b2*-e4i4-MO-injected (4 ng) zebrafish, the pectoral fin length was significantly decreased (to 2.343±0.1634 vs. 4.211±0.1012 X10^2^ μm; P<0.0001 by the unpaired Student’s *t*-test) (Figs. 2, S6, S7).

**Fig. 2.**
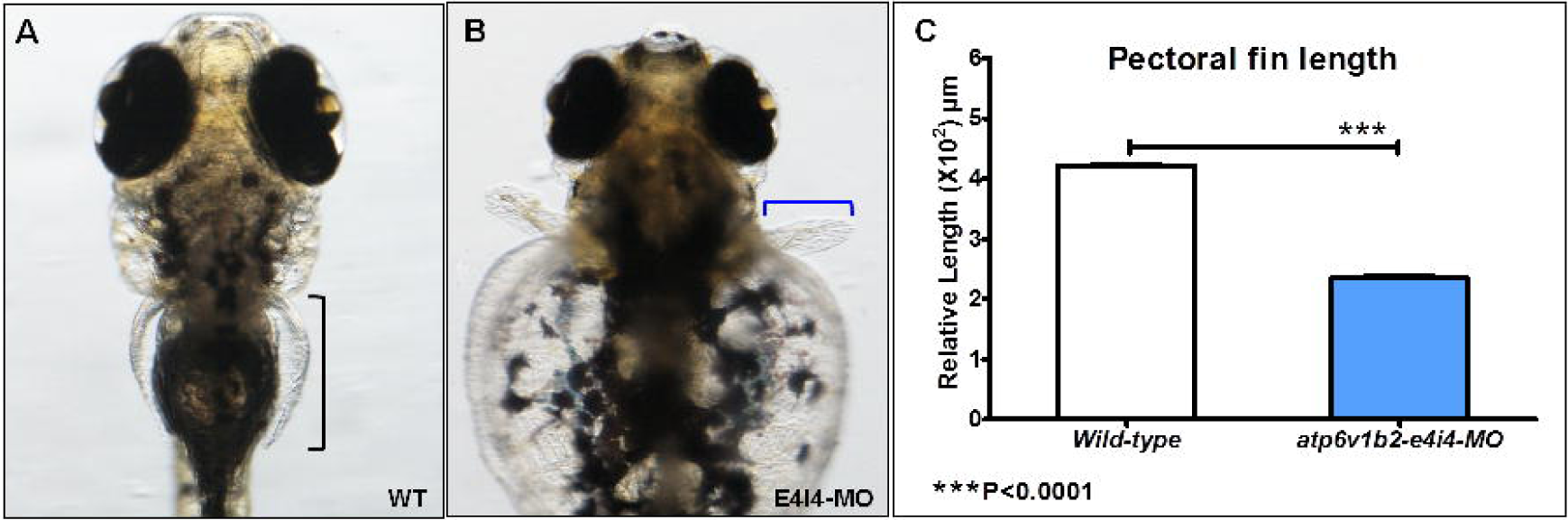
*Atp6v1b2* knockdown disrupts pectoral fin development. (A–B) Ventral views of the pectoral fins of 6 dpf larvae (the fins are bracketed). In uninjected WT controls, the pectoral fins extended from the body and were of normal length at 6 dpf (A, black bracket). In contrast, *atp6v1b2*-e4i4-MO (4 ng) injection reduced pectoral fin growth (B, blue bracket) (Figs. S6, S7). (C) The relative length of the fin fell 1.8-fold in *atp6v1b2* morphants (n = 10) at 6 dpf. (*** P?0.0001 by the unpaired Student’s *t*-test). dpf, days post-fertilisation.

### atp6v1b2 knockdown induces potent hair cell loss

Hair cells of the lateral line can be visualised by simply allowing embryonic zebrafish to swim in dye-containing water. Interneuromast cells are located between the primary neuromasts, and are initially tightly associated with both nerves and glia. These cells migrate ventrally, away from inhibitory nerve cues, and only differentiate into secondary neuromasts (Coffin et al. 2010; Goodrich 2005; Parng et al. 2006; Ton and Parng 2005). Wild-type control embryos and 5 dpf embryos injected with 4 ng *atp6v1b2*-e4i4-MO, 4 ng *atp6v1b2*-ATG-MO, and 8 ng *atp6v1b2*-E13I13-MO were stained with the mitochondrial potentiometric dye DASPEI. Significantly decreased hair cell and head neuromast staining were evident in all three *atp6v1b2* morphants compared to the wild-type control (the average hair cell numbers were 2.40±1.713, 0.80±0.919, and 1.60±1.350; vs. 14.60±0.516; P<0.0001; Figs. 3, S8–S10).

**Fig. 3.**
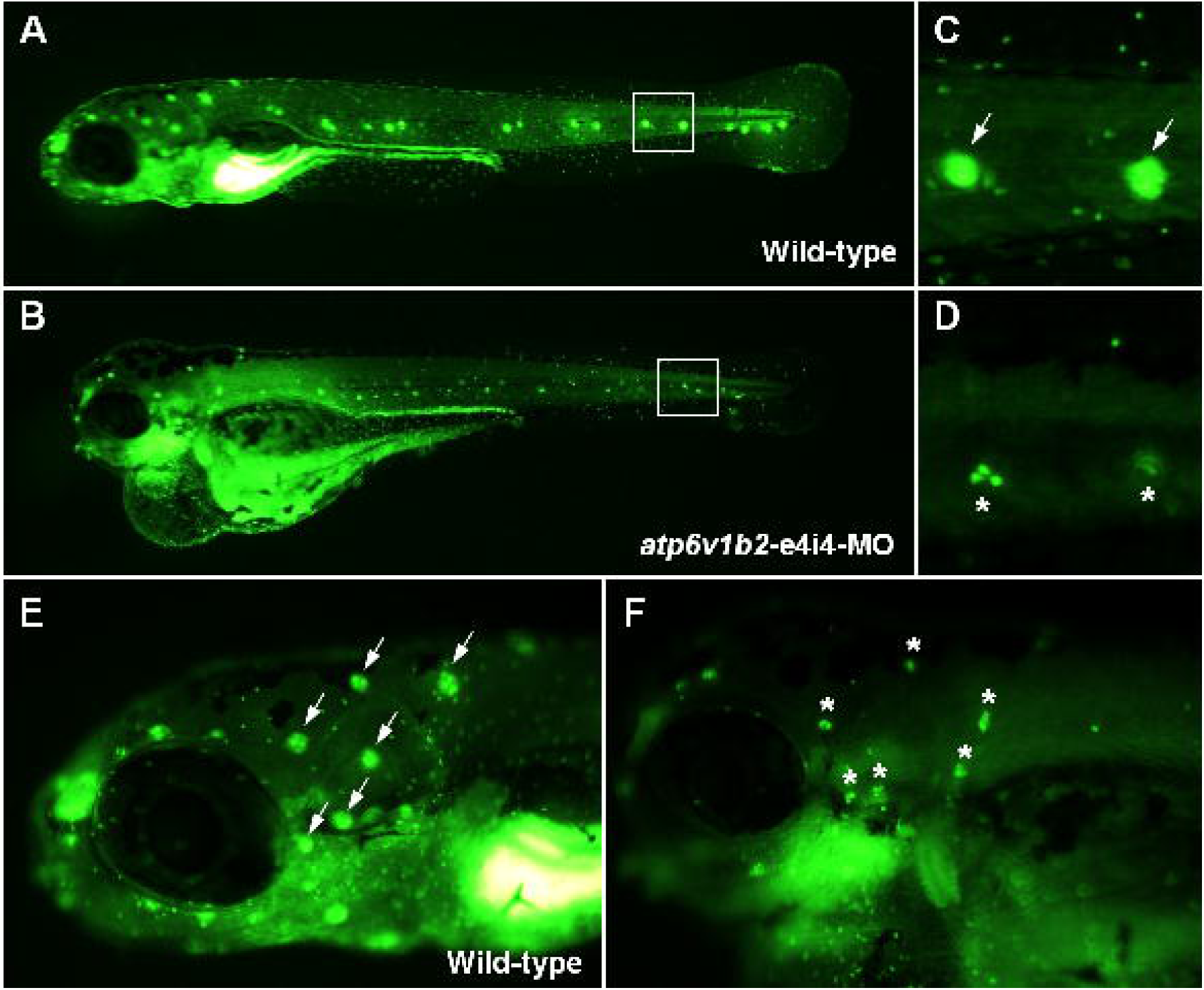
*Atp6v1b2* knockdown induces potent hair cell loss in zebrafish. WT control embryos and embryos injected with 4 ng *atp6v1b2*-e4i4-MO were stained with the mitochondrial potentiometric dye DASPEI at 5 dpf. Hair cells stereotypically located on the lateral line stained as green dots (A, C; white arrows). Uninjected WT zebrafish had normal hair cell numbers. In contrast, significantly reduced hair cell (B, D; asterisks) and head neuromast staining (E, white arrow; F, asterisk) were evident in *atp6v1b2* morphants. The boxed regions are shown at higher magnifications in the right panels. Fluorescent DASPEI images were inverted prior to particle analysis and fluorescence signals from hair cells quantified via morphometry. dpf, days post-fertilisation. Also see Figures S8–S10.

### Effect of atp6v1b2 c. 1516C>T overexpression on the early development of zebrafish

The *de novo* mutation c.1516C>T (p.Arg506X) in *ATP6V1B2* is the genetic etiology of DDOD syndrome in patients (Yuan et al. 2014). To observe the effect of *atp6v1b2* c.1516C>T overexpression on the early development of zebrafish, we injected pcDNA3 *atp6v1b2 c*.*1516C>T* into fertilised one-cell embryos. Mutant atp6v1b2 overexpression caused developmental defects including decreased body length and severe pericardial oedema in 67.12% of embryos (98/146) (Figs. 4, S11). In addition, *atp6v1b2* c.1516C>T overexpression induced potent hair cell loss. Staining of both hair cells and head neuromasts was significantly reduced in *atp6v1b2* c.1516C>T overexpressing zebrafish compared to the wild-type control (the average hair cell numbers were 0.60±0.9661 vs. 14.60±0.516; P<0.0001; Fig. 5).

**Fig. 4.**
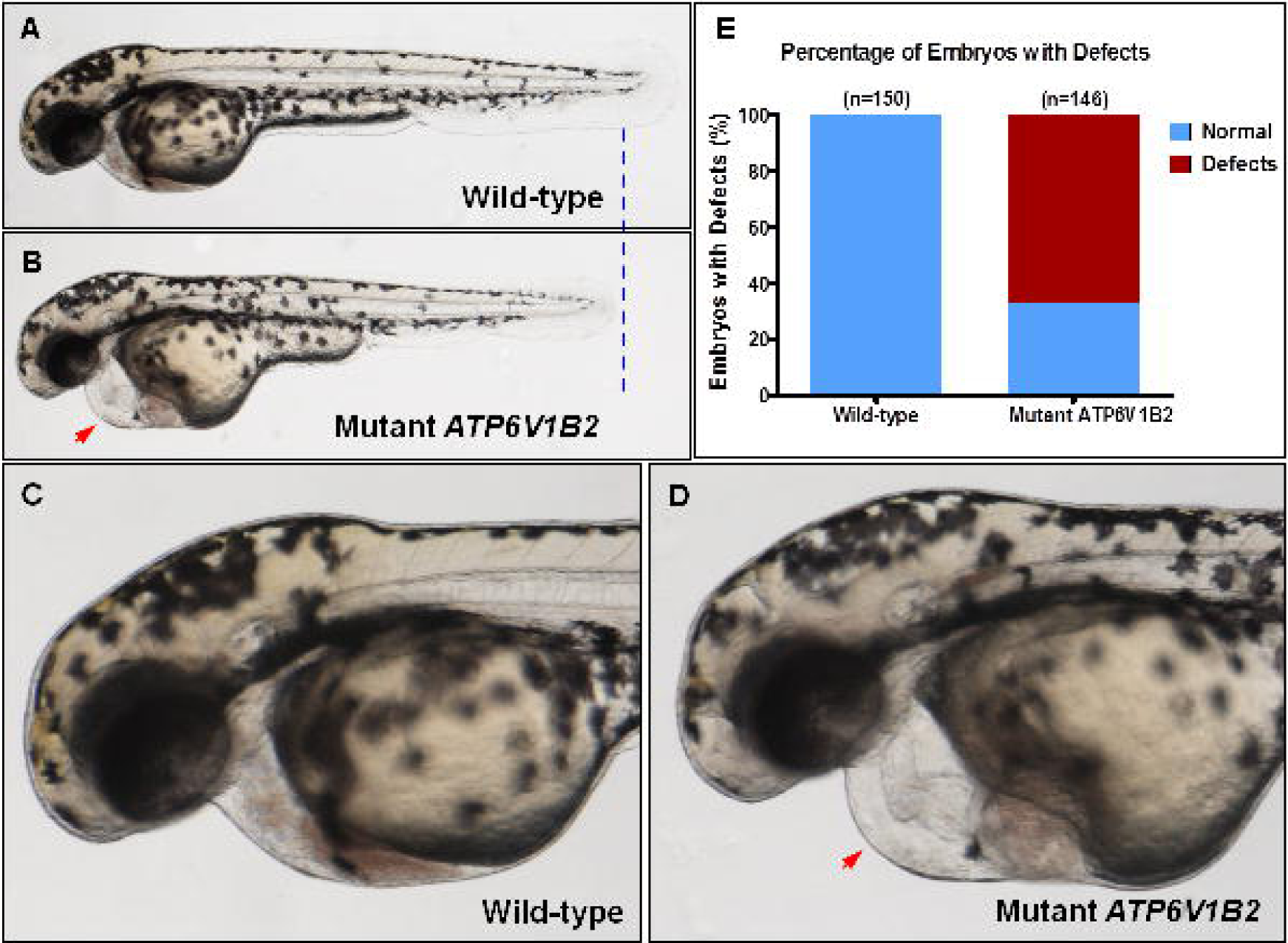
Overexpression of mutant *atp6v1b2* causes developmental defects in zebrafish. Compared to uninjected WT control embryos, mutant human *atp6v1b2* overexpression decreased the body length (blue dotted line) and caused severe pericardial oedema (red arrow) in zebrafish by 52 hpf. The bar graph shows the percentages of embryos with developmental defects (Fig. S11).

**Fig. 5.**
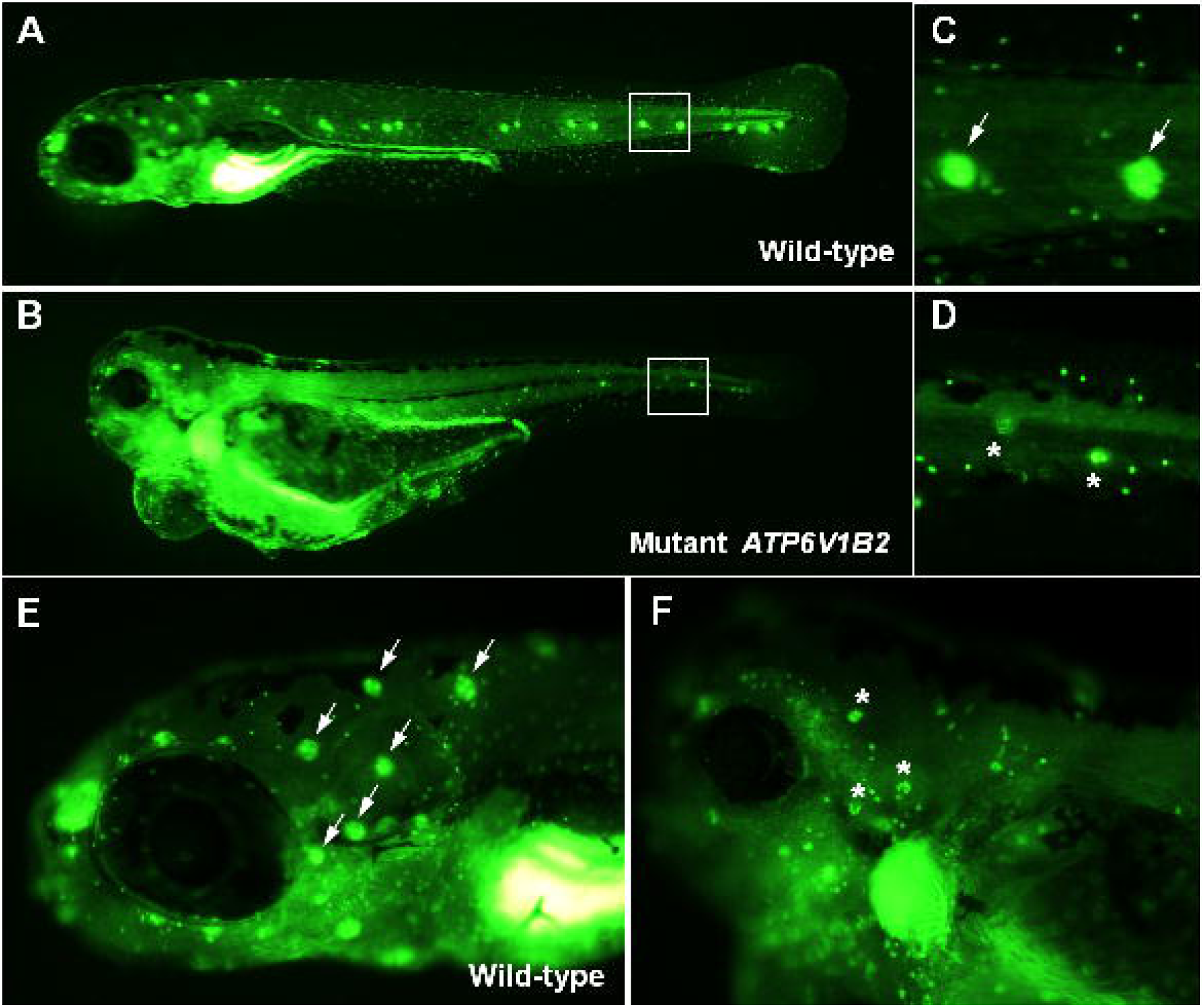
Overexpression of mutant *atp6v1b2* induces potent hair cell loss in zebrafish. WT control embryos and embryos overexpressing mutant *atp6v1b2* were stained with the mitochondrial potentiometric dye DASPEI at 5 dpf. Hair cells stereotypically located on the lateral line stained as green dots (A, C; white arrows). Uninjected WT zebrafish had normal hair cell numbers. In contrast, significantly reduced hair cell (B, D; asterisks) and head neuromast staining (E, white arrow; F, asterisk) was evident in mutant embryos overexpressing *atp6v1b2*. The boxed regions are shown at higher magnifications in the right panels. Fluorescent DASPEI images were inverted prior to particle analysis. Fluorescence signals from hair cells were quantified morphometrically. dpf, days post-fertilisation.

## Discussion

*ATP6V1B2* encodes a component of vacuolar ATPase (V-ATPase, also known as H+-ATPase), which is a multisubunit enzyme that mediates acidification of eukaryotic intracellular organelles. Such acidification is necessary for the efficient operation of various intracellular processes including protein sorting, zymogen activation, receptor-mediated endocytosis, and generation of a proton gradient by synaptic vesicles. V-ATPase is composed of a cytosolic V1 domain and a transmembrane V0 domain. The V1 domain is responsible for ATP hydrolysis, and the V0 domain is responsible for protein translocation. The protein encoded by *ATP6V1B2* is one of the two B-subunit isoforms of the V1 domain, and is highly expressed in organelles of the cerebrum and lysosomes. This V1B2 subunit is usually termed the brain isoform or the lysosomal V1 subunit B2 (Wagner et al. 2004; Nelson et al. 2000). *ATP6V1B2* deficiency has been associated with hereditary diseases such as DDOD syndrome, osteopetrosis, and renal tubular acidosis. It has also been suggested that deficiencies in *ATP6V1B1* and *ATP6V0A4* are associated with distal renal tubular acidosis and hearing loss (Stover et al. 2002; Stehberger et al. 2003; Sobacchi et al. 2001; Sly et al. 1983).

Zebrafish *atp6v1b2*, also termed *vatB1*, shares 92.9% sequence identity with human *ATP6V1B2*. *Atp6v1b2* has a total of 14 exons and encodes a V-ATPase of 509 amino acids. *Atp6v1b2* and its isoform, *atp6v1b1* (also termed *vatB1*), are expressed at all developmental stages of the zebrafish (Schredelseker and Pelster 2004). The two isoforms can be found in various tissues of adult zebrafish, including the gills, heart, liver, spleen, kidney, intestine, swim bladder, and muscle (Boesch et al. 2003). Our finding that *atp6v1b2* is expressed by zebrafish hair cells, further confirms that the protein plays a role in hearing. Although *Atp6v1b2* cochlear knockdown mice suffer hearing loss, whether and how such knockdown triggers onychodystrophy requires further study.

In the present work, we used two types of MOs to knockdown *atp6v1b2* expression. ATG-MO blocks translation, and the other two MOs (E4I4-MO and E13I13-MO) block splicing. The associated phenotypes were decreased body length, severe pericardial oedema, non-inflation of the swim bladder, hair cell loss, and shortened pectoral fins. Zebrafish fins and tetrapod limbs are homologous in the context of early patterning and gene expression (Ito and Handa 2012). Thus, shortening of the pectoral fins in *atp6v1b2*-knockdown zebrafish indirectly supports the suggestion that the gene is involved in the development of tetrapod limbs. E13I13-MO-knockdown zebrafish exhibited a milder phenotype, without any decrease in body length, indicating that atp6v1b2, with more residual amino acids, may also play a role in the early stage of zebrafish body elongation.

The zebrafish otic vesicle begins to form at 16 hpf, and gives rise to the principal sensory components of the inner ear (Haddon and Lewis 1996). Adult zebrafish possess Weberian ossicles connecting the swim bladder to the inner ear. It has been proposed that the inner ear of otophysans can be stimulated by particle motion via a direct pathway, and by sound pressure via an indirect pathway (i.e., the swim bladder and the Weberian ossicles, respectively). The swim bladder of zebrafish starts to inflate at 5 dpf, but Weberian ossicles are absent in zebrafish at less than 1 week of age (Higgs et al. 2003). Therefore, during the first week of development, zebrafish are probably insensitive to sonic pressure because neither the Weberian ossicles nor the swim bladder can deliver pressure stimuli to the inner ear. The saccule and utricle of zebrafish less than 1 week of age are only motion-sensitive to direct stimuli. Larval zebrafish exhibit a startle response to sudden acoustic stimuli at an early stage, thus when free swimming commences. In the present study, the stereotypic escape response was inactive in 48 hpf *atp6v1b2*-e4i4-MO-injected larvae, supporting the notion that impaired atp6v1b2 dulls hearing.

The swim bladder is an air-filled sac dorsally located in the abdominal cavity. The organ allows fish to balance the hydrostatic pressure and reduces the energetic cost of swimming. Morphological and molecular evidence suggests that the swim bladder is evolutionarily homologous to the lung (Zheng et al. 2011; Winata et al. 2009). The endothelia and blood circulation play important roles in the organisation and differentiation of swim bladder structures and swim bladder inflation (Winata et al. 2010). The swim bladder acts as a pressure transducer. *Atp6v1b2*-knockdown morphants exhibited impairment of swim bladder inflation, increasing the energetic cost of swimming, influencing sound pressure conduction, and triggering hearing impairment in adult zebrafish.

DDOD syndrome aside, *ATP6V1B2* mutations are associated with Zimmermann-Laband syndrome (ZLS, MIM 135500) (Kortum et al. 2015), a developmental disorder characterised by facial dysmorphism featuring gingival enlargement, intellectual disability, hypoplasia or aplasia of the nails and terminal phalanges, and hypertrichosis. The DDOD and ZLS syndromes share the phenotype of nail and phalangeal aplasia. The mutations c.1516 C>T of a DDOD family and c.1454 G>C of several ZLS families both arose *de novo*. The *ATP6V1B2* c.1516 C>T (p.Arg506X) mutation segregated with aplasia in three Chinese DDOD families. This nonsense mutation inserts a premature stop codon, yielding a truncated protein lacking the last five amino acids. Mutant *ATP6V1B2* overexpression in zebrafish decreased the body length, caused severe pericardial oedema, and induced potent hair cell loss, but did not affect pectoral fin growth or swim bladder inflation even when the concentration of mutant plasmid attained 200 ng/μL (twice that required for induction of hair cell loss). This can be explained as follows: 1) c.1516 C>T is a dominant haplo-insufficient mutation (Yuan et al. 2014). One copy is inactivated but the other copy is functional. The single functional copy does not produce enough V-ATPase to create wild-type conditions, resulting in dysplasia. We suggest that hair cells of sensory nerves, and the cardiovascular system, are more sensitive to gene and protein defects than the pectoral fin and swim bladder. Alternatively, 2) *atp6v1b2* may be regulated differently in various zebrafish organs during development. Thalidomide induces fin and ear (otic vesicle) defects in zebrafish (Ito et al. 2010; Ito and Handa 2012). However, such teratogenic effects can be partly rescued by overexpression of a functionally active, thalidomide binding-defective, form of zCrbn^YW/AA^ (bearing the p.Y374A and p.W376A mutations). Ito et al. (44) showed that cereblon (CRBN) is a thalidomide-binding protein. CRBN forms a complex with E3 ubiquitin ligase, which in turn damages both DNA binding protein 1 (DDB1) and Cul4A. Both proteins are required for limb outgrowth and expression of fibroblast growth factor Fgf8 in the zebrafish and chicken. Fgf and Bmp signaling act in concert to pattern the cochlea (Puligilla et al. 2007), regulate early cardiogenesis (Alsan and Schultheiss 2002), and control limb bud interdigital programmed cell death (Pajni-Underwood et al. 2007). Whether the functions of *atp6v1b2* during early zebrafish development are mediated by Fgf and/or Bmp signalling requires further investigation.

## Conclusions

We developed an *atp6v1b2* zebrafish knockdown model using three MOs, and observed that body length decreased, severe pericardial oedema developed, the swim bladder was not inflated, hair cells were lost, and the pectoral fins were shortened. This is the first work to show that atp6v1b2 impairment affects early development; the gene plays important roles in the development of hearing, the pectoral fins, the cardiovascular system, and the swim bladder. Such results support the suggestion that the gene is affected in patients with syndromic hearing loss.

## Acknowledgements

These investigations were supported by the National Nature Science Foundation of China (no. 81230020, 81371096), and a grant from the Minister of Science and Technology of China (no. 2012BAI09B02) to P.D. Y.Y.Y. acknowledges grants from the National Nature Science Foundation of China (no. 81371098), the Beijing Natural Science Foundation (no. 7132177), and the Beijing Nova Programme (no. 2009B34). No funder played any role in study design, data collection or analysis, the decision to publish, or preparation of the manuscript.

---

Fig. S1. *Atp6v1b2* knockdown with E4I4-MO causes developmental defects. Compared to WT, *atp6v1b2* knockdown with E4I4-MO decreased body length and caused severe pericardial oedema (red arrow) by 32 hpf and 48 hpf.

Fig. S2. *Atp6v1b2* knockdown with ATG-MO causes developmental defects. Compared to uninjected WT control embryos, embryos injected with 4 ng *atp6v1b2-*ATG-MO (ATG-MO) exhibited a decrease in body length (blue dotted line), severe pericardial oedema (red arrow), and non-inflation of the swim bladder (blue arrow).

Fig. S3. *Atp6v1b2* knockdown with E13I13-MO causes developmental defects in zebrafish. Compared to uninjected WT control embryos, embryos injected with 8 ng *atp6v1b2-*e13i13-MO (E13I13-MO) exhibited severe pericardial oedema (red arrow) and non-inflation of the swim bladder (blue arrow).

Fig. S4. RT-PCR confirming the efficacy of the *atp6v1b2*-e4i4-MO.

A) The zebrafish *atp6v1b2* gene was targeted by a specific antisense MO preventing proper splicing of exon 4 (E4I4-MO). Primers 2F and 5R differentiate WT (non-mutant) transcripts from those in which introns 4–5 have been inserted. Below the diagram, the introns 4–5-inserted transcript of E4I4-MO-injected embryos (404 bp), and that of uninjected embryos (328 bp) are schematically shown.

B) Left: RT-PCR of *atp6v1b2* transcripts from uninjected and E4I4-MO MO-injected embryos at 2 dpf, revealing insertion of introns 4–5. Right: Sanger sequencing of both the WT and introns 4–5-containing band; the sequences were as expected.

Fig. S5. RT-PCR confirming the efficacy of the *atp6v1b2*-e13i13-MO.

A) The zebrafish *atp6v1b2* gene was targeted by a specific MO antisense RNA preventing proper splicing of exon 13 (E13I13-MO). Primers 12F and 14R amplify WT (non-mutant) transcripts and those in which introns 13–14 have been inserted. Below: A schematic depiction of the introns 13–14-insert transcript of E13I13-MO-injected embryos (1,432 bp) compared to that of uninjected embryos (359 bp).

B) Left: RT-PCR of the *atp6v1b2* transcript from uninjected and E13I13-MO MO-injected embryos 2 days after fertilisation, demonstrating insertion of introns 13–14. Right: Sanger sequencing of both the WT and the introns 13–14-containing band; the sequences were as expected.

Fig. S6. *Atp6v1b2* knockdown with E4I4-MO affects pectoral fin development of zebrafish. In WT control fish, the pectoral fins stretched from the body wall and were of normal length by 6 dpf (black bracket). In contrast, *atp6v1b2*-e4i4-MO (4 ng) injection reduced pectoral fin growth (blue bracket)

Fig. S7. *Atp6v1b2* knockdown with ATG-MO or E13I13-MO affects pectoral fin development of zebrafish. Compared to the uninjected WT control, zebrafish injected with 4 ng ATG-MO or 8 ng E13I13-MO were compromised in terms of pectoral fin development (blue bracket).

Fig. S8. *Atp6v1b2* knockdown induces hair cell loss in zebrafish. Each column shows five examples of hair cell loss in the lateral line.

Fig. S9. *Atp6v1b2* knockdown with ATG-MO induces potent hair cell loss in zebrafish. WT 5 dpf control embryos and embryos injected with 4 ng *atp6v1b2*-ATG-MO were stained with the mitochondrial potentiometric dye DASPEI. Hair cells stereotypically located on the lateral line stained as green dots (A, C; white arrow). Uninjected zebrafish had normal hair cell numbers. In contrast, significantly reduced hair cell (B, D; asterisk) and head neuromast staining (F; asterisk) was evident in *atp6v1b2* morphants. The boxed regions are shown at higher magnifications in the right panels. Fluorescent DASPEI images were inverted prior to particle analysis. Fluorescent signals from hair cell were quantified morphometrically. dpf, days post-fertilisation.

Fig. S10. *Atp6v1b2* knockdown with E13I13-MO induces potent hair cell loss in zebrafish. WT 5 dpf control embryos and embryos injected with 8 ng *atp6v1b2*-e13i13-MO were stained with the mitochondrial potentiometric dye DASPEI. Hair cells stereotypically located on the lateral line stained as green dots (A, C; white arrow). Uninjected WT zebrafish had normal hair cell numbers. In contrast, significantly decreased hair cell (B,D; asterisk) and head neuromast staining (F; asterisk) was evident in *atp6v1b2* morphants. The boxed regions are shown at higher magnifications in the right panels. Fluorescent DASPEI images were inverted prior to particle analysis. Fluorescent signals from hairs cell were quantified morphometrically. dpf, days post-fertilisation.

Fig. S11. *Atp6v1b2* c.1516 C>T overexpression causes developmental defects in zebrafish. Body length decreased and severe pericardial oedema (red arrow) developed in zebrafish both 52 hpf and 5 dpf.

The English in this document has been checked by at least two professional editors, both native speakers of English. For a certificate, please see: http://www.textcheck.com/certificate/BJlbef

